# Effects of Single-Session Perturbation-Based Balance Training with Progressive Intensities on Resilience and Dynamic Gait Stability in Healthy Older Adults

**DOI:** 10.1101/2025.01.05.631356

**Authors:** Ringo Tang-Long Zhu, Friederike A. Schulte, Navrag B. Singh, Christina Zong-Hao Ma, Chris Awai Easthope, Deepak K. Ravi

## Abstract

Single-session perturbation-based balance training (PBT) has demonstrated improvements in dynamic stability during the initial step following perturbation in older adults. However, its broader effects on comprehensive balance recovery remain inconclusive. This pilot randomized controlled trial investigated the impact of personalized single-session PBT on reactive balance control during walking, employing advanced stability analysis techniques. Ten participants in the training group underwent a single session consisting of 32 unpredictable treadmill-induced slips and trips of progressively increasing intensity, while ten participants in the control group engaged in unperturbed treadmill walking. Key outcome measures included margin of stability (MoS) parameters: minimum MoS and the number of recovery steps, and resilience parameters: peak instability and recovery time, assessed at baseline, immediately post-intervention, and three months post-intervention following an unexpected treadmill slip. The training group exhibited significant immediate and sustained improvements (p < 0.05) in minimum MoS values, alongside a notable reduction in peak instability (p < 0.05) immediately post-intervention. These changes were not observed in the control group. However, neither group demonstrated significant alterations in the number of recovery steps or recovery time across the assessment periods. In conclusion, single-session PBT enhanced reactive balance control by improving the magnitude of post-perturbation responses, but it did not significantly influence the speed of recovery to baseline conditions.

## 1 Introduction

Falls and fall-related injuries are common health problems among older adults^1–4^ and carry a significant societal cost^5,6^. Balance and functional exercises (e.g., sit-to-stand, stepping) can reduce fall incidence in community-dwelling older adults by around 24%^7^, and are a key part of recommended guidelines for fall prevention^8^. However, these exercises primarily target older adults’ volitional balance control and have less focus on the reactive balance control^9^. Volitional balance control involves intentional and feedforward movements to maintain stability. In contrast, reactive balance control involves automatic responses to unpredictable perturbations and relies on feedback mechanisms. This type of control is crucial for quickly recovering from slips or trips, the leading cause of falls during outdoor activities in community-dwelling older adults^10^. Over the past two decades, perturbation-based balance training (PBT) has emerged as a promising fall-prevention method designed to enhance reactive balance control^11^. Through PBT, individuals learn to react and adapt to sudden losses of balance, gradually developing effective balance recovery strategies^11^. Importantly, PBT aligns with recent guidelines recommending balance-challenging exercises to prevent falls in community-dwelling older adults, emphasizing the need for more high-quality evidence^8^.

Traditionally, PBT is conducted over multiple sessions and has been shown to enhance reactive balance control and reducing fall incidence by 22% to 44%^12,13^. However, multi-session programs can be time-consuming and may suffer from low adherence rates^14^. Recent studies have explored whether delivering multiple perturbations within a single session could offer similar benefits. One study reported a 50% reduction in fall incidence following single-session PBT^14^, suggesting this approach may reduce the need for prolonged exercise and improve engagement. Research on single-session PBT has primarily focused on two key outcomes. In terms of real-life falls, a single session of 24 overground slips significantly reduced falls over one year^14^, whereas a single session of 40 treadmill slips did not lower the 6-month fall incidence^9^. Regarding dynamic stability, studies often use the “margin of stability” (MoS)^15–22^ to assess the effects of single-session PBT on reactive balance control, typically during the first recovery step after an unexpected perturbation^15–20^. A greater positive MoS reflects improved gait stability^23^, with both immediate^16–18,20–22^ and long-term^15,19^ benefits observed in several studies. In summary, while single-session PBT improves dynamic stability, its impact on real-life fall reduction remains inconsistent. This may be attributed to small sample sizes, as fall incidence studies require large-scale trials for reliable results. Variations in training intensity and duration may also play a role^9,14^. Additionally, analyzing only the first recovery step might overlook important aspects of fall prevention, suggesting the need for broader assessments in future research.

Exploration of the optimal training dose and personalized protocols for single-session PBT has been limited. Training doses play a critical role - doses that are too low may be ineffective in reducing fall risk, while doses that are too high could lower adherence and may not provide additional benefits in fall prevention beyond a certain threshold^24^. Most previous studies have implemented a fixed number of perturbations, constant perturbation intensity, and standardized scenarios for single-session PBT^9,14–16,19–22^. Only two studies have investigated how the dose of single-session PBT influences dynamic stability and reactive balance control^17,18^. Lee et al. found that 40 treadmill slips did not produce better immediate outcomes compared to 24 slips^18^. In contrast, Wang et al. demonstrated that single-session PBT with progressively increasing intensities led to better immediate outcomes than a control group performing unperturbed treadmill walking^17^. Furthermore, few studies have incorporated enriched perturbation scenarios (such as a combination of slips and trips) within single-session PBT. Tailoring the training dose and incorporating diverse perturbation scenarios could provide a more comprehensive training experience, potentially enhancing fall prevention outcomes.

Another research gap is that analyzing the immediate response of dynamic stability after a perturbation may not fully capture the entire balance recovery and fall-avoidance mechanism. Measuring the minimum MoS following an unpredictable perturbation provides critical insight into the most unstable state during recovery^25^. Additionally, calculating the number of steps needed to return to baseline MoS helps assess recovery speed^25,26^. Several advanced non-linear analysis techniques have also been proposed to evaluate balance control performance. Unlike traditional linear methods (e.g., standard deviation), which interpret variability around the mean as random noise, non-linear analysis suggests that fluctuations in normal human walking reflect patterns indicative of complex adaptive behavior^27^. Ravi et al. introduced a method to assess steady-state behavior by leveraging the natural variability in walking patterns^28^. This approach also evaluates resilience - defined as the ability to resist perturbations or recover to steady state following a disturbance^28,29^. Two key indicators of resilience can be derived: peak instability, defined as the maximum deviation from steady-state behavior following a perturbation, and recovery time, defined as the duration from peak instability to the return to steady state^28^. By evaluating both the magnitude of the response (e.g., minimum MoS, minimum MoS relative to baseline, peak instability) and the speed of recovery (e.g., number of recovery steps, recovery time), we can gain deeper insights into the effects and mechanisms underlying single-session PBT. This approach can also help identify key factors that influence the success or failure of this type of training.

Given the identified research gaps, the primary objective of this study was to investigate the immediate and retained effects of single-session PBT on reactive balance control in older adults. The training incorporated enriched perturbation scenarios, featuring both treadmill slips and trips, with progressive intensities organized in blocks. Resilience and anteroposterior MoS parameters were evaluated following an unpredictable treadmill slip, comparing older adults who received the single-session PBT to those who did not, across different time points (i.e., pre-intervention, post-intervention, and three months post-intervention). It was hypothesized that: 1) Single-session PBT would result in greater reductions in peak instability, recovery time, and the number of recovery steps, along with greater increases in minimum MoS values and minimum MoS values relative to baseline from pre- to post-intervention, compared to the control group. 2) These improvements would persist at three months post-intervention compared to pre-intervention. The secondary objective was to determine whether single-session PBT could enhance balance performance, as measured by clinical tests, and reduce the incidence of prospective falls in older adults. It was hypothesized that the training group would show better clinical test results and experience fewer falls following the single-session PBT compared to the control group.

## 2 Methods

### 2.1 Study Design and Participants

This pilot assessor-blinded, parallel-group randomized controlled trial (RCT) was conducted at ETH Zurich, Switzerland, from June 2022 to June 2023. The trial protocol obtained ethical approval from the ETH ethics committee (Ethics Number: 2021-N-90). The flow of intervention and assessment procedures is displayed in **Figure 1**. The Consolidated Standards of Reporting Trials (CONSORT) guidelines were followed.

**Figure 1.**
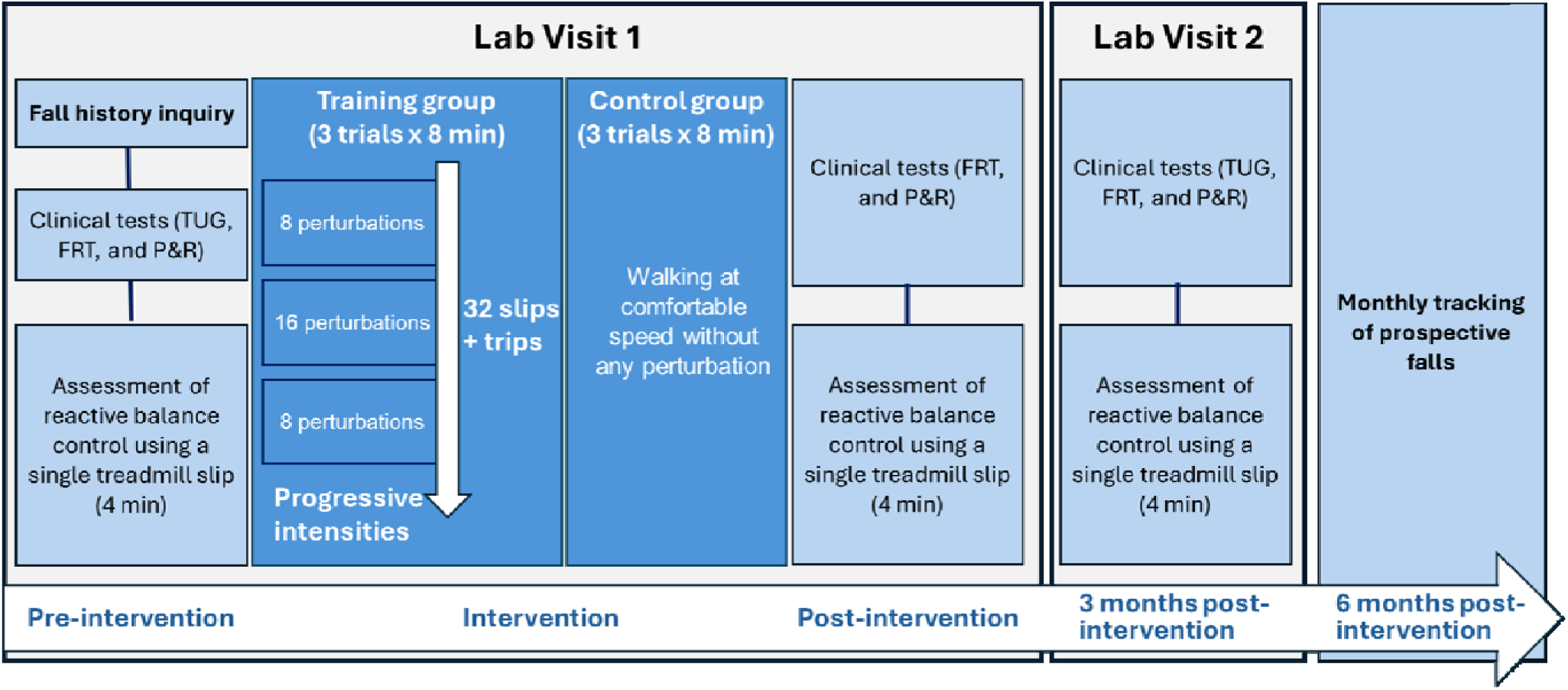
Flowchart of intervention and assessment procedures. TUG: timed up and go test. FRT: functional reach test. P&R: push and release test.

Participants were recruited through convenience sampling using flyers and mail at local physiotherapy centers, the local ergotherapy association, the general practitioner association, and the ETH pensioner’s association. The inclusion criterion was community-dwelling people aged 65 years or older. The exclusion criteria were: a) diagnosed with acute or chronic musculoskeletal or neurological impairments, b) having undergone any lower limb surgery such as joint replacement, within the past 1.5 years, c) regular intake of antidepressants, and d) inability to walk on a treadmill for 30 minutes. All participants provided written informed consent. The participants were randomly allocated to the training or control group in a 1:1 ratio. Demographic data, including each participant’s age, sex, height, weight, leg dominance, and fall history within past one year were collected.

### 2.2 Equipment and Intervention

A commercially available treadmill with an overhead safety harness (Quasar, H/P/Cosmos Sports & Medical GmbH, Nussdorf-Traunstein, Germany) was used for treadmill walking. A 12-camera motion capture system (Vicon T160, Vicon Motion Systems, Oxford, United Kingdom), sampling at 200 Hz, was used for kinematic data collection. A total of 62 reflective markers were affixed to the participant’s whole-body anatomical landmarks (see **Supplementary Table 1**). The treadmill was synchronized with the motion capture system to induce unpredictable perturbations. Specifically, a customized MATLAB script identified heel strikes (defined as the minimal vertical foot velocity per gait cycle) in real time^30^. The heel strike data were then transmitted to Visual Studio, where a custom script sent a trigger to the treadmill, causing it to accelerate/decelerate the belt immediately following a detected heel strike. The perturbation occurred during the single-stance phase.

Prior to the intervention, the maximum treadmill belt accelerations that challenged each participant’s forward and backward limits of standing stability without requiring a step were determined by gradually increasing the treadmill belt acceleration in 0.25 m/s² increments. The participant’s self-selected comfortable walking speed was determined for both the control and intervention groups by gradually increasing the treadmill belt speed in 0.2 km/h increments. The final speed was calculated as the average of the speeds at which the participant reported feeling slightly too fast and slightly too slow.

#### 2.2.1 Personalized Single-session PBT (Training Group)

Each participant completed a single session of PBT during treadmill walking at a comfortable speed. The session consisted of three eight-minute trials, with two-minute intervals for rest between trials. The perturbations included equal numbers of treadmill trips (belt acceleration) and slips (belt deceleration), randomly delivered to either foot. Each participant experienced 8 unpredictable perturbations in the first trial, 16 in the second trial, and another 8 in the third trial. The intensity of the perturbations progressively increased across the trials. In the first trial, the perturbation intensities of treadmill trips and slips were set at 1.25 times the previously determined forward and backward limits of standing stability, respectively. If the participant felt comfortable with the intensity and no fall occurred, the perturbation intensity was increased by 0.5 m/s^2^ for the next trial.

#### 2.2.2 Unperturbed Walking (Control Group)

Each participant walked on the treadmill at a comfortable speed without any perturbations. This also comprised three eight-minute trials, with two-minute rest intervals between trials.

### 2.3 Primary Outcome Measures

Each participant’s reactive balance control during treadmill walking was assessed pre-intervention, immediately post-intervention, and three months post-intervention (**Figure 1**). During each assessment, the participant walked on the treadmill for four minutes, and a single treadmill slip (2.25 m/s² belt deceleration) was randomly induced to the participant’s dominant leg. The participant was instructed not to talk, scratch, or adjust clothes during the treadmill walking, while the whole-body kinematics were collected. The primary outcome measures, namely, MoS and resilience, were processed and analyzed to quantify each participant’s reactive balance control performance during treadmill walking.

Before the calculation of MoS and resilience, the three-dimensional coordinates of all reflective markers were first low-pass filtered (Butterworth, zero-phase lag, fourth-order, 5Hz cut-off frequency). The sacrum marker was then used as an approximate representation of CoM^31^.

#### 2.3.1 MoS

The MoS in the anteroposterior direction was calculated at each foot strike using the equations provided below ^32,33^. The heel strikes identified in real time during treadmill walking were initially used as foot strikes^30^. If the participant did not land on the heel following an unpredictable perturbation, motion capture data was checked in Vicon Nexus software to manually confirm the foot strike time points.

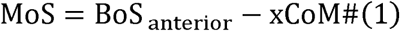

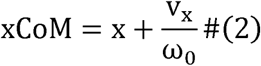

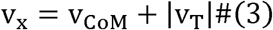

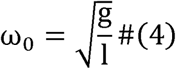

Briefly, the anteroposterior MoS was the difference between the anterior boundary of the base of support, BoS _anterior,_ and the extrapolated CoM position, xCoM (**Figure 2A**). The BoS _anterior_ was defined as the position of the toe marker of the leading leg in the anteroposterior direction^25^. The xCoM represented the anteroposterior extrapolated CoM position. Here, x was anteroposterior position of the CoM, and v_x_ was anteroposterior CoM velocity relative to the treadmill belt, which equaled the sum of the anteroposterior CoM velocity relative to ground and the absolute value of treadmill belt velocity. The ω_0_ was eigenfrequency of pendulum, the g was gravitational acceleration, and l was pendulum length which equaled the distance between CoM position and midpoint of the line connecting medial malleolus marker and lateral malleolus marker^34^.

**Figure 2.**
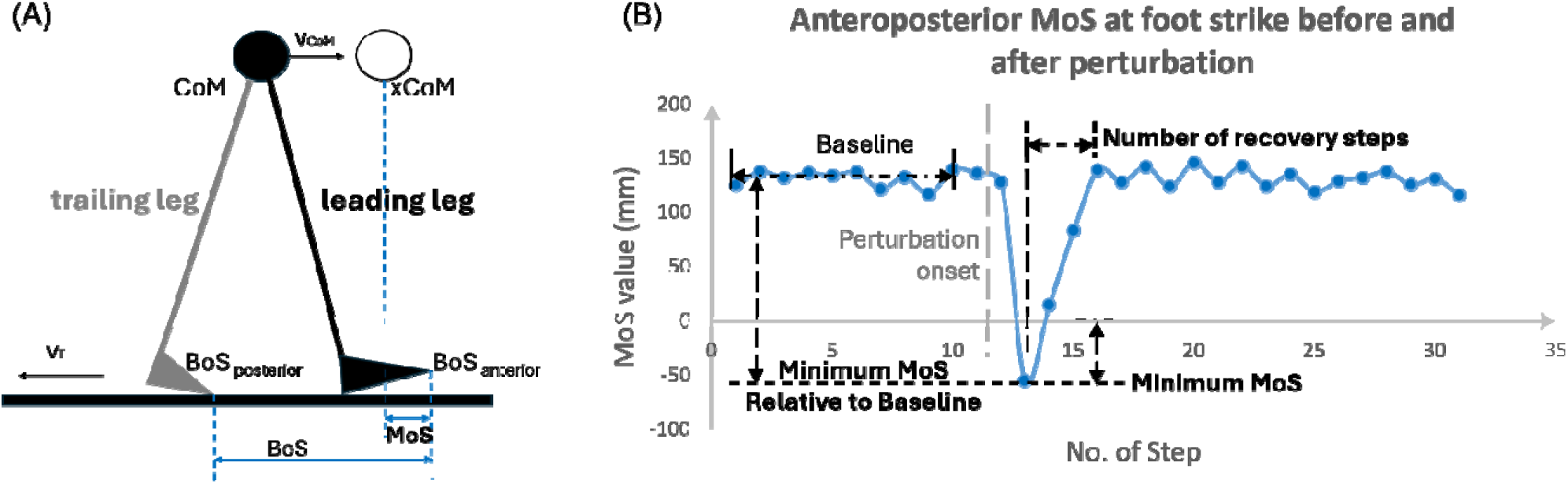
Calculation and analysis of margin of stability (MoS). (A) Schematic illustration of anteroposterior MoS during treadmill walking. CoM: center-of-mass position. xCoM: extrapolated center-of-mass position. v_CoM_: center-of-mass velocity relative to ground. v_T_: treadmill belt velocity relative to ground. BoS _anterior_: toe marker position of leading leg. BoS _posterior_: toe marker position of trailing leg. (B) Illustration of analyzed parameters of anteroposterior MoS during treadmill walking.

MoS values at foot strikes for the 10 consecutive steps before and 20 steps after the perturbed step were analyzed (**Figure 2B**). The baseline MoS was defined as the mean of the MoS values from the 10 steps prior to the perturbed step. The minimum MoS was obtained to represent the most unstable post-perturbation gait relative to zero^25^. Its positive sign indicated a stable gait, meaning the xCoM was within the BoS^23^. The minimum MoS relative to baseline was further determined by taking the minimum MoS minus the baseline MoS, dividing by the baseline MoS, and multiplying by 100%. It represented the most unstable post-perturbation gait relative to pre-perturbation walking. The number of recovery steps was calculated as the number of steps from the one with the minimum MoS to the first step where the MoS returned to within one standard deviation (SD) below the baseline MoS^26^.

#### 2.3.2 Resilience

A participant’s resilience in reactive balance control was quantified by performing state space reconstruction of vertical CoM time series to determine the pre-perturbation steady state boundaries and evaluate the post-perturbation CoM response^28^. The processing procedures are detailed below.

The pre-perturbation and post-perturbation vertical CoM time series were reconstructed separately in state space using the time-delay embedding procedure^35^ which involved the determination of a time lag τ and an embedding dimension d. The resulting state space vectors [X(t), X(t+τ), …, X(t+(d-1)∗τ)] were further reduced to three dimensions [X(t), X(t+τ), X(t+2∗τ)]. The mean of all vectors on the reconstructed pre-perturbation trajectory was taken as the centroid and a reference trajectory (M) was fitted to these vectors using an eight-term Fourier model. An ellipse was constructed around M at each integer angle ranging from 0° to 359°. The semi-major and semi-minor axes of the ellipse were set to twice the largest and second-largest SDs, respectively, calculated from the 50 nearest vectors in three dimensions. All ellipses together formed a torus (T_2σ_), representing the boundaries of steady-state behavior (**Figure 3A**). Then the Euclidean distance, D(t), from each vector on the reconstructed post-perturbation trajectory to M was calculated (**Figure 3B**). The peak instability, defined as the maximum Euclidean distance, was analyzed to indicate the most unstable state following an unpredictable perturbation. The recovery time was defined as the time interval from the peak instability point to the recovery point, which was identified when the trajectory re-entered the torus (T_2σ_) and remained within it for three consecutive gait cycles, allowing for up to five outliers. Both this criterion and a more stringent criterion (allowing one outlier for five consecutive gait cycles) were tested. The current criterion was ultimately chosen, as it produced more reasonable recovery time values (see **Supplementary Table 2**).

**Figure 3.**
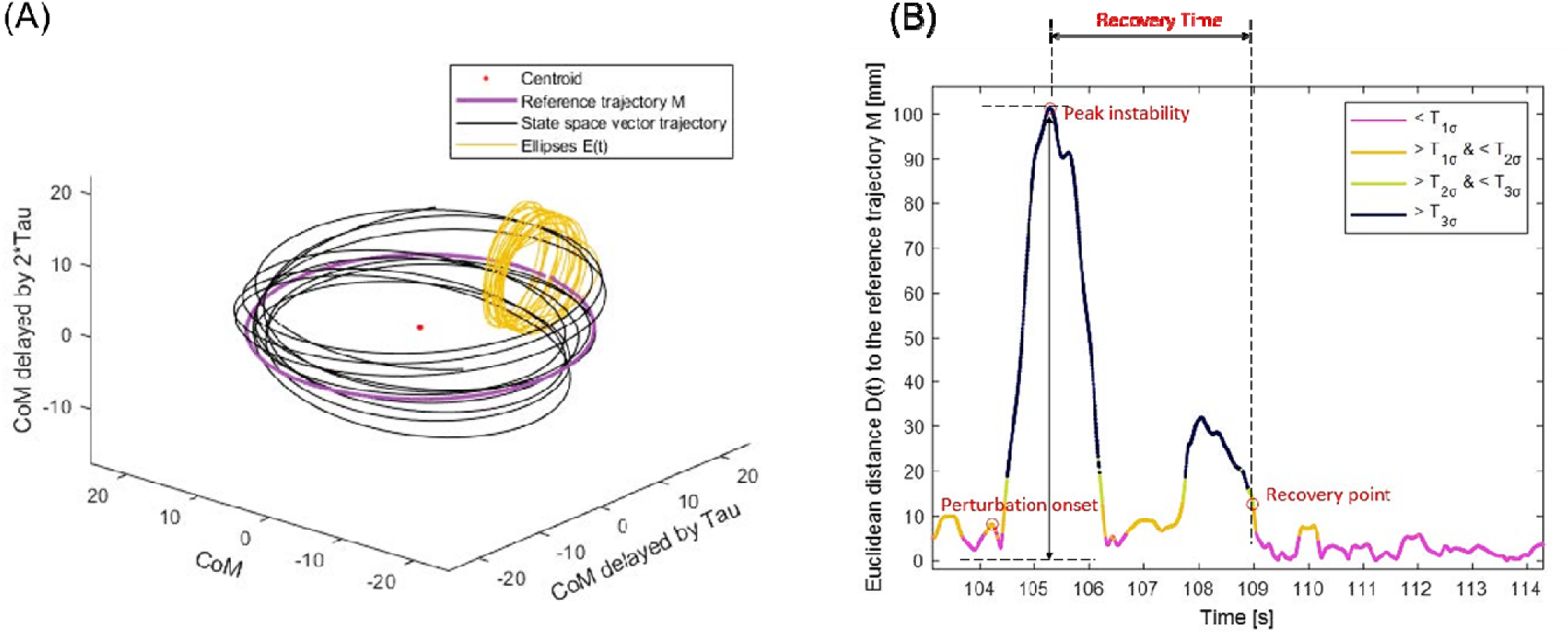
Processing and analysis of resilience. (A) Illustration of using state space reconstruction of vertical center of mass (CoM) displacement time series to obtain steady state. The ellipses E around the reference trajectory M indicate the boundaries for steady-state behavior. (B) Euclidean distances of state space vectors from the reconstructed post-perturbation trajectory to the reference trajectory M.

### 2.4 Secondary Outcome Measures

Clinical tests were conducted to assess each participant’s postural balance performance (**Figure 1**). The Timed Up and Go (TUG) test was performed pre-intervention and again three months post-intervention to evaluate volitional balance control. Participants were instructed to “stand up from the chair, walk a 3-m distance, walk back, and sit down,” with the completion time recorded^36^. The Functional Reach Test (FRT) was conducted pre-intervention, immediately post-intervention, and three months post-intervention to evaluate volitional balance control. Participants were instructed to “raise the arm, make a fist, and reach forward as far as possible without making a step,” with the reached distance measured^37^. The Push and Release (P&R) test was administered pre-intervention, immediately post-intervention, and three months post-intervention to evaluate reactive balance control. Participants were instructed to “lean backward against the examiner’s hands placed on their scapula”; the examiner then suddenly removed their hands to induce a stepping response, and the participant’s performance was rated using a specific scale^38^. The long completion time of TUG test, small reaching distance of FRT, and high score of P&R test indicate poor balance performance.

Monthly phone calls were conducted to follow up each participant’s number of prospective falls over the 6 months following the intervention (**Figure 1**). A fall was defined as an unintentional event that led to the person coming to the ground or a lower level^5^.

### 2.5 Statistical Analyses

Data of demographics and prospective falls were compared in training vs. control groups as below. For continuous variables (i.e., age, body mass index [BMI], number of previous falls, and number of prospective falls), independent sample t-tests were used when data were normally distributed, while Mann-Whitney U tests were used for when data were not normally distributed. For dichotomous categorical variables (i.e., sex, dominant leg, fall history status, and prospective fall status), Fisher’s exact tests were used considering the small sample size.

Effects of “intervention” (between-subjects factor: training group vs. control group) and “time” (within-subjects factor: pre-, immediately post-, and three months post-intervention) on the outcomes of postural balance performance were examined as below. For each continuous outcome (i.e., the completion time of TUG test, the reached distance in FRT, the minimum MoS, the minimum MoS relative to baseline, the number of recovery steps, the peak stability, or the recovery time), a two-way mixed analysis of variance (ANOVA) followed by post hoc pairwise comparisons with Bonferroni corrections was conducted. The interaction effect of “intervention” and “time”, the main effect of each factor, and the simple effect of each factor were examined. Effect sizes (*f*) of interaction effects for primary outcome measures were calculated. The *f* values of 0.1, 0.25, and 0.4 represented small, medium, and large effect sizes, respectively^39^. For the ordinal outcome (i.e., the score of P&R test), a Mann-Whitney U test was used to compare group difference at each assessment, and a Friedman test followed by post hoc pairwise comparisons with Bonferroni corrections was conducted to examine changes over time within each group.

Post hoc correlation analyses were conducted between clinical test outcomes and primary outcomes for each time point of assessment. A Pearson correlation test was used if the two continuous variables followed normal distributions; otherwise, a Spearman correlation test was used. All statistical analyses were conducted in SPSS version 25.0 with the two-tailed significance level set at 0.05.

## 3 Results

Twenty-two eligible older adults were recruited for this study initially. One participant was excluded due to technical issues, and another one participant was excluded due to developing a secondary health problem. In the end, 20 participants were included in the data analysis, with 10 in the training group and 10 in the control group. As presented in **Table 1**, participants in the training group were significantly younger than those in the control group by five years (*p <* 0.01). There were no significant differences between groups in body mass index (BMI), sex, leg dominance, fall history status, or the number of previous falls (*p >* 0.05).

**Table 1.** Results of demographics, prospective falls, and clinical tests.

|  | **Training group**<br>**(n = 10)** | **Control group**<br>**(n = 10)** |
| --- | --- | --- |
| **Age (year)** | 67.1 ± 2.8* | 72.8 ± 5.2* |
| **BMI (kg/m<sup>2</sup>)** | 23.6 ± 3.0 | 22.8 ± 2.4 |
| **Female (number, percentage)** | 5, 50% | 5, 50% |
| **Right leg dominance (number, percentage)** | 8, 80% | 10, 100% |
| **Fall history status** |  |  |
| Fallers (number, percentage) | 2, 20% | 5, 50% |
| Recurrent fallers (number, percentage) | 1, 10% | 4, 40% |
| **Number of previous falls** | 0 (0) | 1 (2) |
| **Prospective fall status** |  | **n = 8** |
| Prospective fallers (number, percentage) | 3, 30% | 3, 37.5% |
| Prospective multiple fallers (number, percentage) | 2, 20% | 1, 12.5% |
| **Number of prospective falls** | 0 (1.0) | 0 (1.0) |
| **TUG test (s)** |  |  |
| Pre-intervention | 8.2 ± 1.4 | 8.6 ± 1.4 |
| Three months post-intervention | 8.2 ± 1.1 | 9.3 ± 2.1 |
| **FRT (cm)** |  |  |
| Pre-intervention | 30.8 ± 3.6<sup>#</sup> | 28.7 ± 6.9 |
| Immediately post-intervention | 29.5 ± 6.3<sup>†</sup> | 26.1 ± 6.6 |
| Three months post-intervention | 38.7 ± 4.5<sup>#†*</sup> | 31.5 ± 7.0* |
| **P&R test (score)** |  |  |
| Pre-intervention | 0 (1.0) | 1.0 (1.0) |
| Immediately post-intervention | 0.5 (1.0) | 1.0 (2.0) |
| Three months post-intervention | 0.5 (1.0) | 1.0 (0) |

Notes: Normally distributed continuous data were displayed as mean ± standard deviation. Non-normally distributed continuous data or ordinal data were displayed as median (interquartile range). BMI: body mass index. Fallers: people with ≥ 1 previous fall. Recurrent fallers: people with ≥ 2 previous falls. Prospective fallers: people with ≥ 1 prospective fall. Prospective multiple fallers: people with ≥ 2 prospective falls. TUG: timed up and go. FRT: functional reach test. P&R: push and release.
\* represents the significant difference between training group and control group ( $p < 0.05$ ). # or † represents the significant change over time, and the same symbols represent the pairwise comparison with significant difference ( $p < 0.05$ ).

### 3.1 MoS

One participant in the control group held the handrails of treadmill for balance recovery following the unpredictable perturbations during assessment trials. This participant was excluded from the MoS analysis (**Figure 4**), as the calculation of MoS assumes the absence of external forces other than gravity and ground reaction forces^33^.

**Figure 4.**
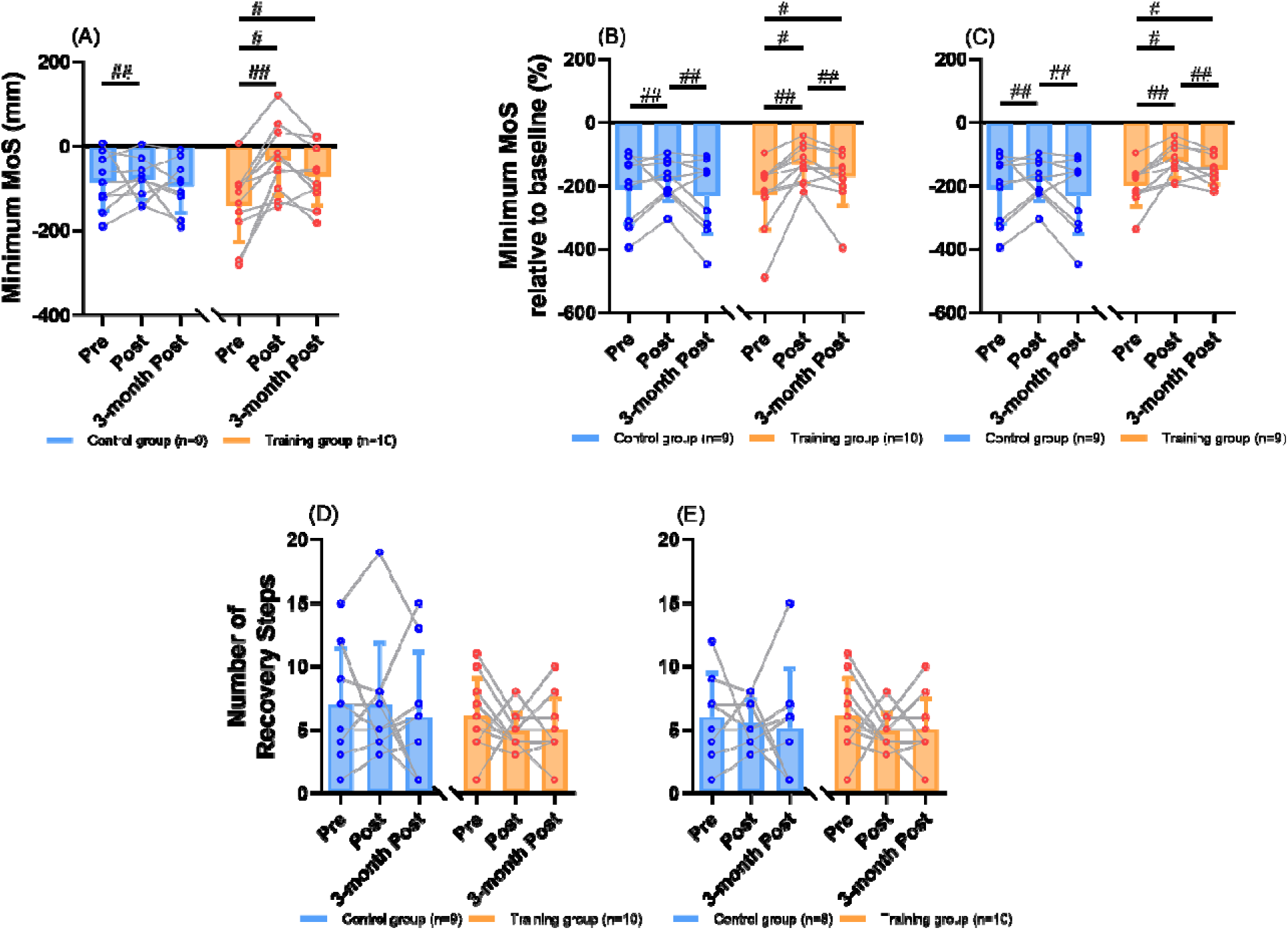
The MoS outcomes (mean ± SD) with individual values for comparisons between groups and among time points. (A) Results of the minimum MoS. (B) Results of the minimum MoS relative to baseline. (C) Results of the minimum MoS relative to baseline after excluding the extreme value. (D) Results of the number of recovery steps. (E) Results of the number of recovery steps after excluding the extreme value. # represents the significant change over time in a certain group (simple effect of “time”, *p <* 0.05). MoS: margin of stability. SD: standard deviation.

For the minimum MoS, there was a significant main effect of “time” (**Figure 4A**), showing that participants of two groups had generally increased minimum MoS values immediately post-intervention than pre-intervention (*p <* 0.01). There was no significant main effect of “intervention” (*p* > 0.05). There was a significant interaction effect of “intervention” and “time” (*p <* 0.01; *f* = 0.64). Significant simple effect of “time” existed within the training group (**Figure 4A**), where participants’ minimum MoS values significantly increased immediately post-intervention (-33 ± 84 mm [mean ± SD; same as below]; *p <* 0.001) and three months post-intervention (-71 ± 70 mm; *p <* 0.01) as compared to pre-intervention (-140 ± 87 mm). No significant simple effect of “time” was observed within the control group (pre-intervention: -85 ± 67 mm, immediately post-intervention: -78 ± 48 mm, three-month post-intervention: -94 ± 62 mm; *p >* 0.05). There was no significant simple effect of “intervention” at any time point (*p >* 0.05).

For the minimum MoS relative to baseline, there was a significant main effect of “time” (**Figure 4B**), showing that participants of two groups had generally larger values immediately post-intervention than pre-intervention (*p <* 0.01) or three-months post-intervention (*p =* 0.028). There was no significant main effect of “intervention” (*p >* 0.05). There was a significant interaction effect of “intervention” and “time” (*p* = 0.035; *f* = 0.47). Significant simple effect of “time” existed within the training group (**Figure 4B**), where participants’ minimum MoS values relative to baseline significantly increased immediately post-intervention (-131% ± 58%; *p <* 0.001) and three months post-intervention (-172% ± 90%; *p <* 0.01) as compared to pre-intervention (-227% ± 111%). No significant simple effect of “time” was observed within the control group (pre-intervention: -210% ± 109%, immediately post-intervention: -182% ± 65%, three-month post-intervention: -231% ± 119%; *p >* 0.05). There was no significant simple effect of “intervention” at any time point (*p >* 0.05). The identified significant results remained after excluding one participant in the training group who had extremely small minimum MoS relative to baseline (**Figure 4C**).

For the number of recovery steps, there were no significant main effects (*p >* 0.05), interaction effect (*p >* 0.05; *f* = 0.14), or simple effects of “intervention” and “time” (**Figure 4D**; *p >* 0.05). The training group had recovery steps of 6 ± 3, 5 ± 1, and 5 ± 2, while the control group had 7 ± 4, 7 ± 5, and 6 ± 5 at pre-intervention, immediately post-intervention, and three months post-intervention, respectively (**Figure 4D**). No significant results were found, either, after excluding one participant in the control group who responded to the unpredictable perturbations by running and did not return to baseline BoS within 20 post-perturbation steps (**Figure 4E**; *p >* 0.05).

### 3.2 Resilience

For the peak instability, there was a significant main effect of “time” (**Figure 5A**), showing that participants of two groups had generally lower peak instability values immediately post-intervention than pre-intervention (*p =* 0.014) or three-months post-intervention (*p =* 0.017). There was no significant main effect of “intervention” (*p =* 0.086). No significant interaction effect of “intervention” and “time” was observed (*p >* 0.05; *f* = 0.29). The training group had peak instability values of 57 ± 25, 34 ± 14, and 45 ± 14 mm, while the control group had 60 ± 18, 49 ± 17, and 64 ±

**Figure 5.**
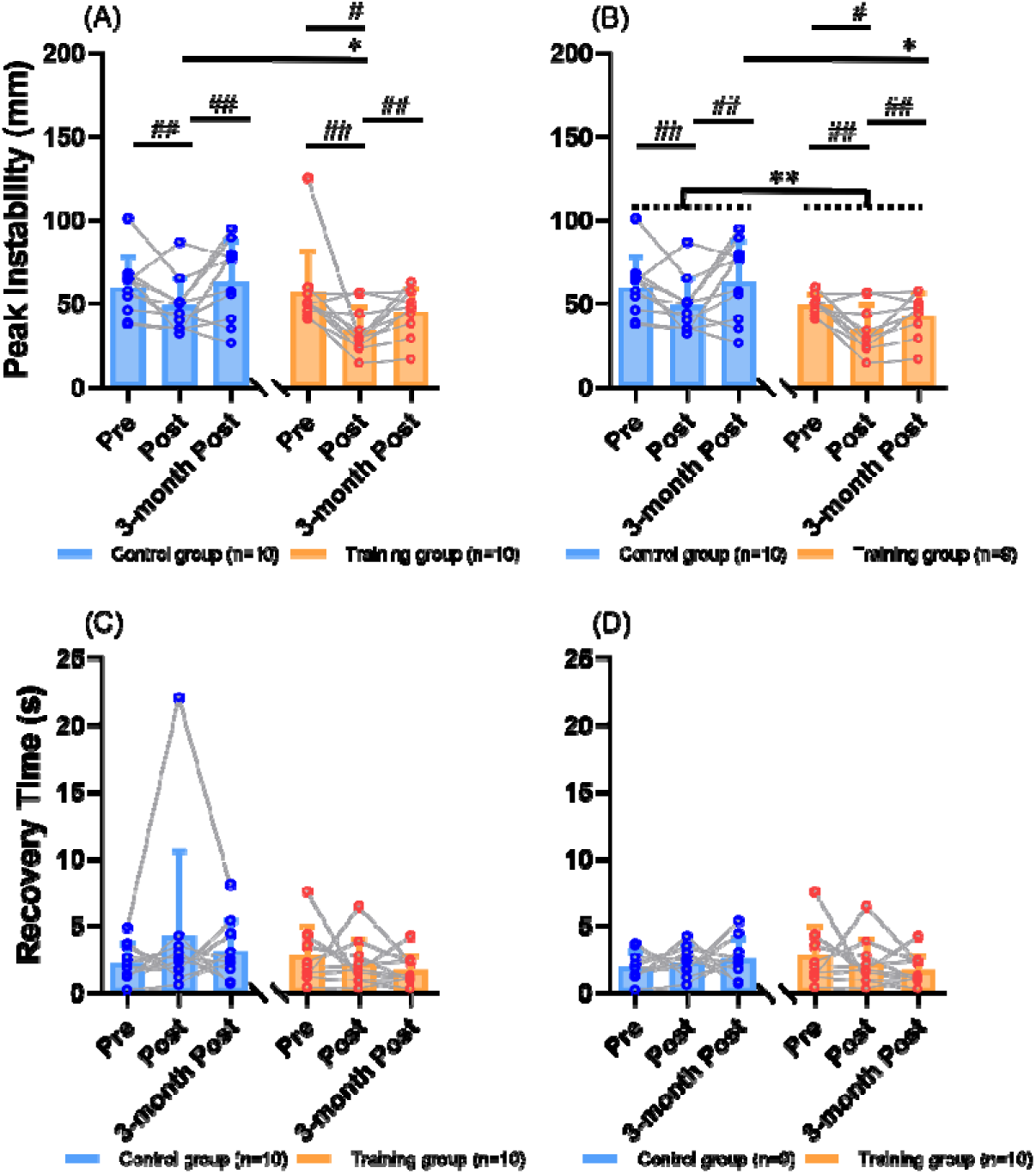
The resilience outcomes (mean ± SD) with individual values for comparisons between groups and among time points. (A) Results of the peak instability. (B) Results of the peak instability after excluding the extreme value. (C) Results of the recovery time. (D) Results of the recovery time after excluding the extreme value. # represents the significant change over time in a certain group (simple effect of “time”, *p <* 0.05). * represents the significant difference between training group and control group at a certain time point (simple effect of “time”, *p <* 0.05). SD: standard deviation.

24 mm at pre-intervention, immediately post-intervention, and three months post-intervention, respectively. Significant simple effect of “time” existed within the training group, where the peak instability values significantly decreased from pre-intervention to immediately post-intervention (*p =* 0.019, **Figure 5A**). No significant simple effect of “time” was observed within the control group (*p =* 0.078). Significant simple effect of “intervention” existed at immediately post-intervention, where participants in the training group showed significantly lower peak instability values than the control group (*p =* 0.044, **Figure 5A**). These significant results were still observed after excluding one participant in the training group who showed extremely high peak instability, except for the changes below (**Figure 5B**). Significant main effect of “intervention” was observed, showing that the training group had generally lower peak instability values than the control group for all time points (*p =* 0.027, **Figure 5B**). Significant simple effect of “intervention” existed at three months post-intervention, where participants in the training group showed significantly lower peak instability values than the control group (**Figure 5B**, *p =* 0.038).

For the recovery time, there were no significant main effects (*p >* 0.05), interaction effect (*p >* 0.05; *f* = 0.28), or simple effects of “intervention” and “time” (**Figure 5C**; *p >* 0.05). The training group had recovery time of 2.8 ± 2.2, 2.2 ± 1.8, and 1.7 ± 1.2 s, while the control group had 2.2 ± 1.4, 4.3 ± 6.3, and 3.1 ± 2.3 s at pre-intervention, immediately post-intervention, and three months post-intervention, respectively (**Figure 5C**). No significant results were found, either, after excluding the participant in the control group who responded by running following unpredictable perturbations and showed an extremely long time to return to steady state (**Figure 5D**; *p >* 0.05).

### 3.3 Clinical Test Results and Prospective Falls

Clinical test results are summarized in **Table 1**. For the FRT, participants in the training group showed a significantly larger reaching distance three months post-intervention than pre-intervention or immediately post-intervention (*p <* 0.01), which was not observed in the control group. Additionally, participants in the training group showed a significantly larger reaching distance than those in the control group at three-month post intervention (*p =* 0.014). For the completion time of TUG test and the P&R score, no significant differences were observed between groups or across time points (*p >* 0.05).

The correlations between clinical test outcomes and primary outcomes were mostly non-significant (**Table 2**). At pre-intervention, the longer completion time of TUG test was significantly related to higher minimum MoS (*r* = 0.596, *p* < 0.01) and higher minimum MoS relative to baseline following an unpredictable perturbation (ρ = 0.574, *p* = 0.01), and the smaller reaching distance of FRT was significantly related to fewer recovery steps following an unpredictable perturbation (*r* = 0.574, *p* = 0.01). At three months post-intervention, the larger score of P&R test was significantly related to the lower minimum MoS relative to baseline (ρ = -0.490, *p* = 0.033) and longer recovery time (ρ = 0.491, *p* = 0.028) following an unpredictable perturbation.

**Table 2.**
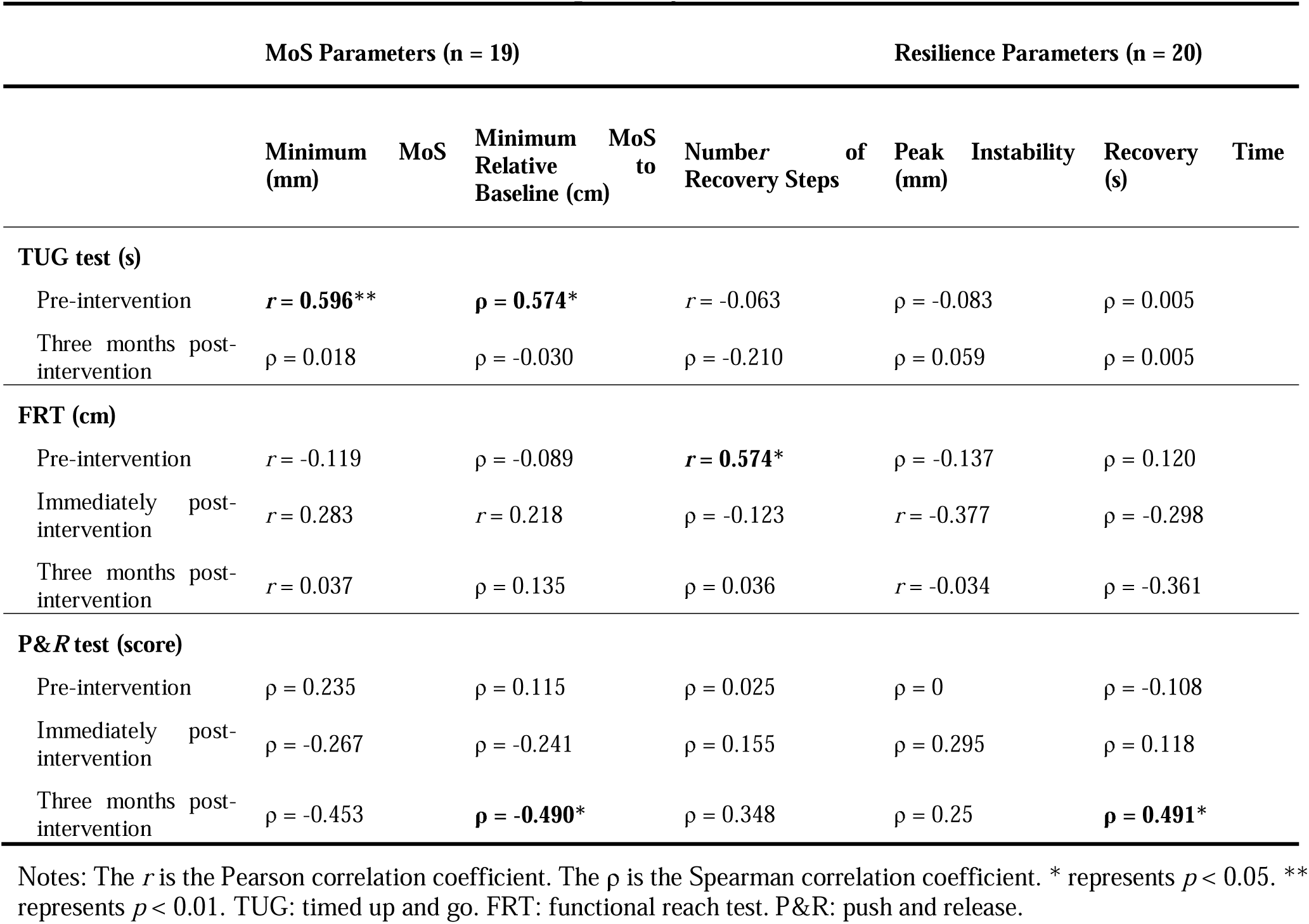
Correlation test results between primary outcomes and clinical test outcomes.

During the tracking of prospective falls, two participants in the control group were lost to follow-up. No significant differences were found in prospective fall status or the number of prospective falls between the two groups (**Table 1**, *p >* 0.05).

## 4 Discussion

The primary objective of this pilot RCT was to quantify how a single-session PBT, involving mixed slips and trips of progressive intensities, affected older adults’ reactive balance performance during walking. Contrary to our hypotheses, the personalized single-session PBT did not lead to improvements across all examined outcomes. It enhanced older adults’ reaction magnitudes (i.e., minimum MoS, minimum MoS relative to baseline, peak instability) in response to unpredictable perturbations. However, it did not significantly improve their recovery speed to pre-perturbation levels (i.e., number of recovery steps, recovery time).

Large effect sizes (f > 0.4) suggest that the single-session PBT had both immediate and lasting effects in improving minimum MoS values and their baseline-relative measures, as well as an immediate effect in reducing peak instability following unpredictable treadmill slips in older adults. The initial slip led to a reduction in step length, contributing to a lower MoS during the first few steps after the perturbation. This trend is consistent with previously reported stride-to-stride MoS responses following a perturbation ^25,40^. The improved minimum MoS parameters indicate that the single-session PBT enhanced dynamic stability during the most unstable phase of reactive balance control. This finding aligns with previous studies showing that single-session PBT increased dynamic stability during the first post-perturbation step^17,19,21^, although few studies have examined minimum MoS in stride-to-stride responses. Additionally, this study found that the single-session PBT reduced the peak deviation of the CoM from the steady state immediately after the intervention. Collectively, these findings on reaction magnitudes suggest that a single-session PBT can enhance older adults’ stability during the most unstable moments of gait or CoM displacement in reactive balance control.

The single-session PBT did not significantly reduce the number of recovery steps or the recovery time following unpredictable treadmill slips. Notably, previous studies have not directly shown that PBT accelerates balance recovery to pre-perturbation levels. Rather, they have highlighted participants’ adaptations in balance recovery speed^25,40^ and muscle activation speed^41,42^. As previously reported, young adults^25^ and older adults without a fall history^40^ were faster (i.e., taking fewer recovery steps to attain baseline MoS) in response to later unpredictable trips than to earlier ones. This discrepancy may be partly due to the greater number of (32 versus 8 perturbations^25,40^) experienced in this study, which may have led to increased fatigue among participants, hindering their adaptation in recovery steps to baseline MoS and recovery time to return to CoM steady state. Further investigation into how participants adapted to perturbations during the single-session training is warranted to optimize the appropriate dose of PBT. For example, if the number of recovery steps or recovery time plateaus as perturbation number or intensity increases, this may indicate that the dose is sufficient for the participant.

Although the single-session PBT produced similar responses in MoS and resilience parameters, it is important to note that these measures reflect different aspects of balance control. Both MoS and resilience parameters captured improvements in reaction magnitudes but did not show changes in recovery speed following unpredictable perturbations. However, there are key differences between the two measures that warrant further consideration. First, MoS represents how far the extrapolated horizontal displacement of the CoM is from the boundary of the BoS^32^, whereas resilience is derived from the reconstruction of vertical CoM displacements and assesses how far the post-perturbation state deviates from the steady state^28^. Second, MoS is typically calculated at specific time events (e.g., foot strike), representing instantaneous dynamic stability^15–22^. In contrast, resilience is derived from state space reconstruction of adjacent vertical CoM displacement time series, capturing the variability of vertical CoM displacements over time^28^. These differences may explain the varying effect sizes observed between MoS parameters (minimum MoS and baseline-relative MoS: large; number of recovery steps: small) and resilience parameters (peak instability: medium; recovery time: small) in quantifying the effects of the single-session PBT. The suspensory strategy, which involves lowering the CoM height, has been recognized as a key balance control mechanism^42–44^. Resilience parameters offer an additional nonlinear assessment of vertical CoM displacement, complementing prior linear analyses. When combined with MoS parameters, resilience measures may provide a more comprehensive evaluation of balance performance and the effects of training.

Contrary to our hypotheses for the secondary objective, not all clinical test results indicated improvements, and there was no reduction in prospective falls following the personalized single-session PBT. The clinical test results suggest that the single-session PBT primarily enhanced the limits of volitional standing stability (as measured by the FRT), rather than influencing the number of reactive steps (P&R test) or volitional gait speed (TUG test). These results align with previous research demonstrating the effects of PBT on TUG test performance^45–49^. Although the impact of PBT on FRT performance has been reported in post-stroke individuals^50^ and those with knee osteoarthritis^51^, evidence for its effectiveness in community-dwelling older adults remains limited. The clinical test results from this study further reflect the findings related to reaction magnitudes and recovery speed following unpredictable perturbations during walking, reinforcing the impact of single-session PBT on postural balance performance. However, the single-session PBT did not lead to a reduction in prospective falls among older adults over the 6-month follow-up period. A previous study similarly found that a single session of treadmill slips in 70 older adults did not reduce prospective falls over a 6-month period^9^, while another study reported that a single session of overground slips in 67 older adults led to a reduction in falls over 12 months^14^. The primary reason for these discrepancies may be the duration of fall tracking, as prospective falls tend to increase exponentially with aging and the accumulation of risk factors^52^. Additionally, factors such as training dose and sample size may have influenced the effectiveness of single-session treadmill-based PBT in preventing falls.

The MoS and resilience parameters may serve as valuable complements to existing tests for assessing balance control and fall risk in older adults. While prospective fall incidence remains the most direct measure of PBT effectiveness in fall prevention, tracking falls over time can be labour-intensive and slow. Clinical tests, such as the TUG test - where a completion time exceeding 15 seconds indicates fall risk^8^ - offer a quick way to identify at-risk individuals. However, these tests may lack the sensitivity needed to detect subtle changes in balance control, particularly in well-functioning community-dwelling older adults, like those in the current study. Moreover, clinical test outcomes showed mostly non-significant correlations with MoS and resilience parameters, suggesting that these measures assess different aspects of balance control. By supplementing clinical tests, MoS and resilience parameters can provide a more comprehensive evaluation of balance performance and the effects of single-session PBT.

The current study has several limitations. First, the small sample size may have limited the ability to detect significant training effects from the single-session PBT. For example, although changes in recovery time after the intervention were not statistically significant, they showed a medium effect size (f = 0.28). Based on this effect size, a future study would require an estimated total of 36 older participants to achieve sufficient power (0.95) for detecting significant changes in recovery time using a two-way mixed ANOVA design (two groups × three time points) at α = 0.05. Second, while the training and control groups were matched for fall history, sex ratio, BMI, and other factors, their ages were not matched in this pilot study. This mismatch may have introduced confounding effects when assessing the impact of the single-session PBT. Although an analysis of covariance (ANCOVA) with age as a covariate was considered, the data did not meet the necessary assumptions. Future RCTs may benefit from stratified randomization, categorizing participants into age groups before randomly assigning them to the training or control groups within each stratum. Third, discrepancies exist between the calculation of MoS in this study and the recommended methods^33^. Specifically, toe markers from both feet were used to define the anteroposterior boundaries of the BoS, as the treadmill used in this study could not record center-of-pressure (CoP) boundaries, as recommended^33^. Additionally, the sacrum marker was used as a proxy for the CoM to ensure comparability with resilience calculations^28,29^, although ideally, the CoM would be estimated using a whole-body kinematic model^33^. These limitations should be carefully considered when interpreting the findings of this pilot study.

In conclusion, this pilot RCT demonstrated that a single session of 32 unpredictable treadmill slips and trips at progressively increasing intensities enhanced reaction magnitudes by improving dynamic stability during the most unstable phase of gait and reducing CoM deviation from the steady state during reactive balance control. However, it did not significantly improve the speed of recovery to unperturbed walking. Future studies with larger, age-matched samples in both training and control groups are necessary to confirm the effects of single-session PBT on both reaction and recovery following unpredictable balance perturbations during walking.

## 5 Data availability

The original contributions presented in the study are included in the article and supplementary material, further inquiries can be directed to the last author.

## 7 Acknowledgments

This research was funded by the Swiss Innovation Agency (grant number: **52519.1 INNO-LS**). We would like to thank Leonie Wunderli, Raphael Schweri, and Olivia Groth who performed experiments and initial data analysis of this study. We would like to also thank Angela Frautschi, Livia Rätzo, Yong Kuk Kim, Mirko Kaiser, and Björn Zimmermann for their help in experiments. We want to thank Fengyi Wang for her help interpret the results. We want to finally thank all the older participants in this study.

## 8 Author contributions

RZ: Formal analysis, Methodology, Software, Visualization, Writing – original draft; FS: Investigation, Resources, Writing – review & editing; NS: Funding acquisition, Writing – review & editing; WT: Resources, Supervision, Writing – review & editing; CM: Resources, Supervision, Writing – review & editing; CE: Conceptualization, Methodology, Resources, Writing – review & editing; DR: Conceptualization, Data curation, Investigation, Methodology, Project administration, Resources, Software, Visualization, Supervision, Writing – review & editing.

---

**Supplementary Table 1.**
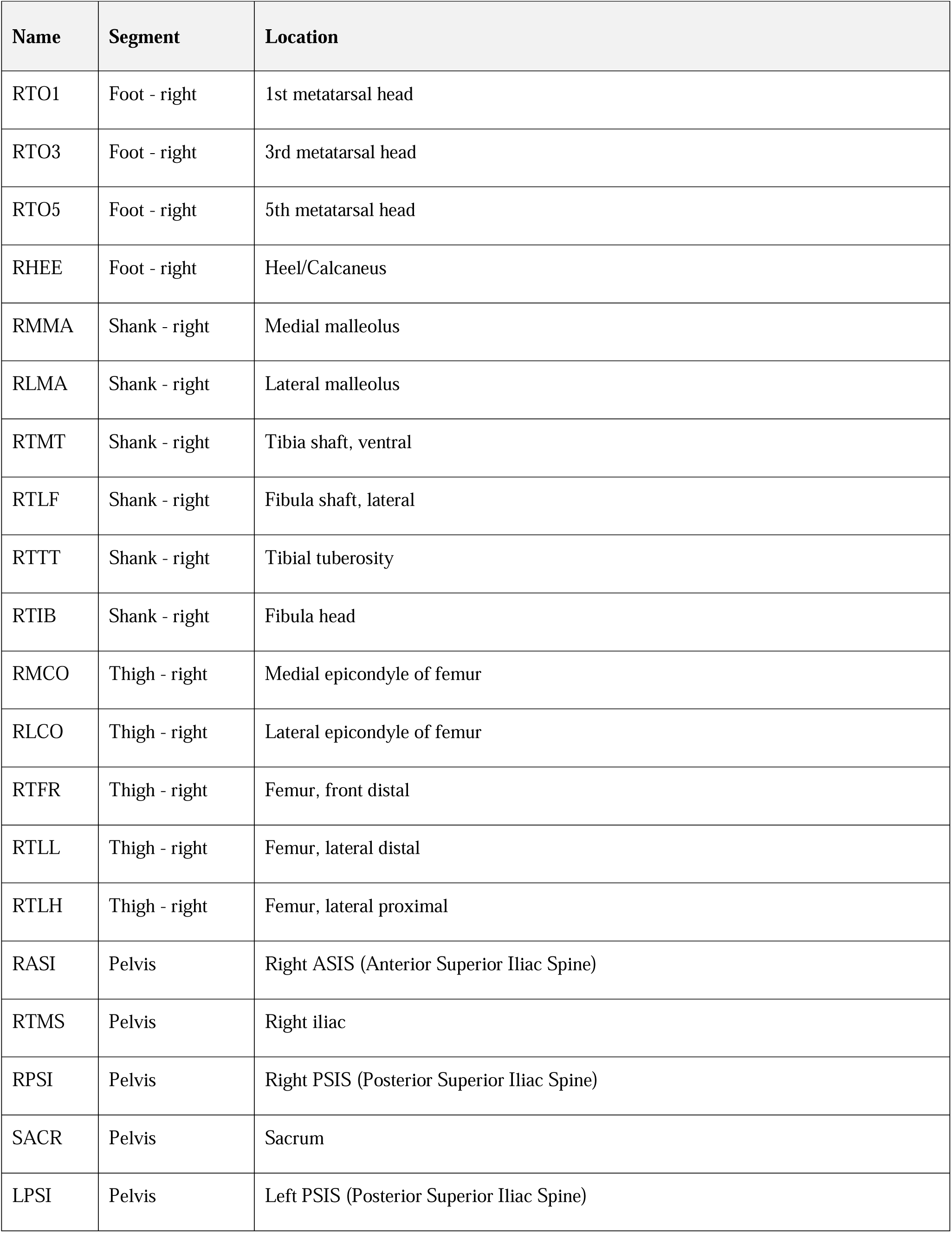
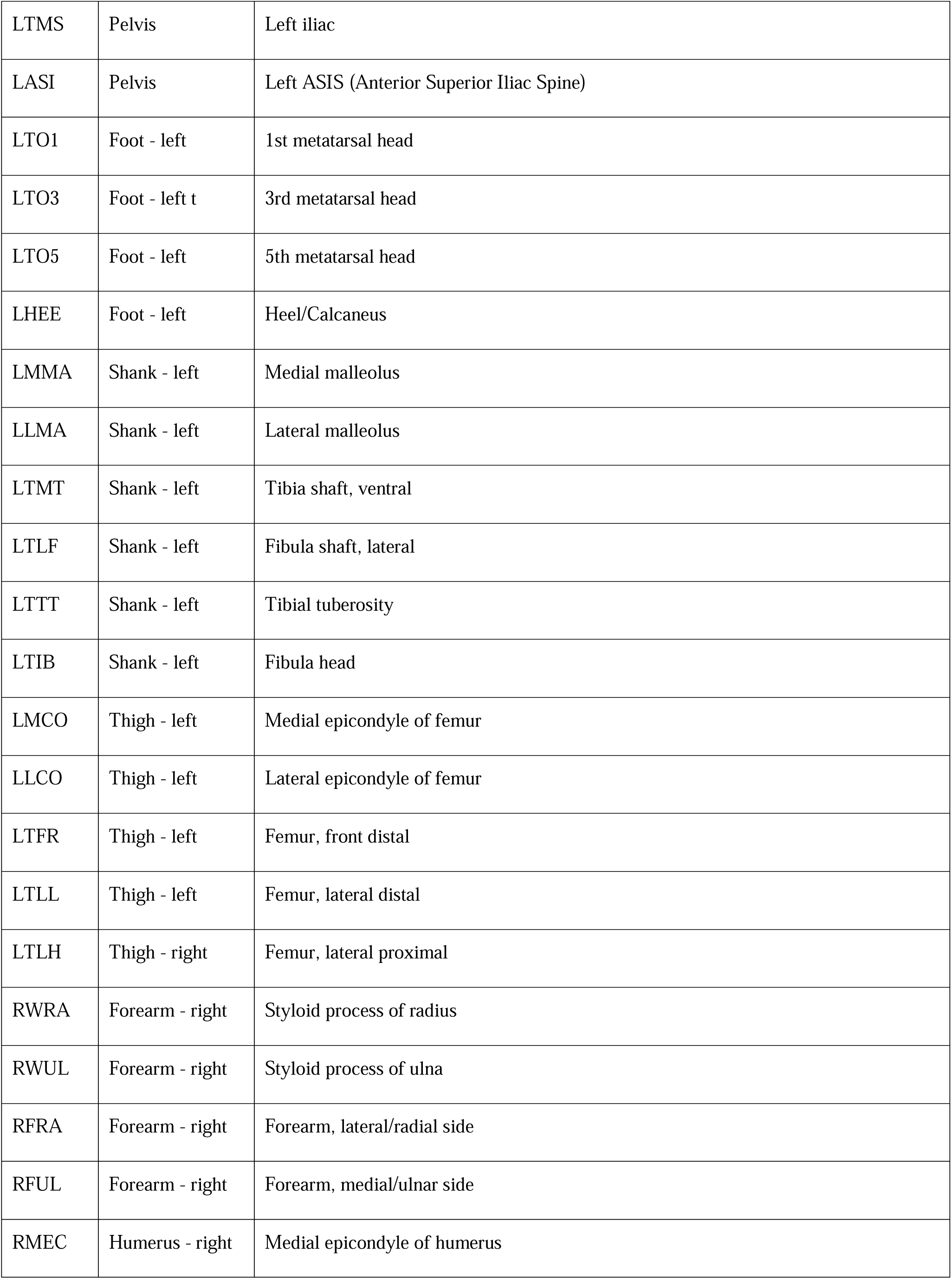
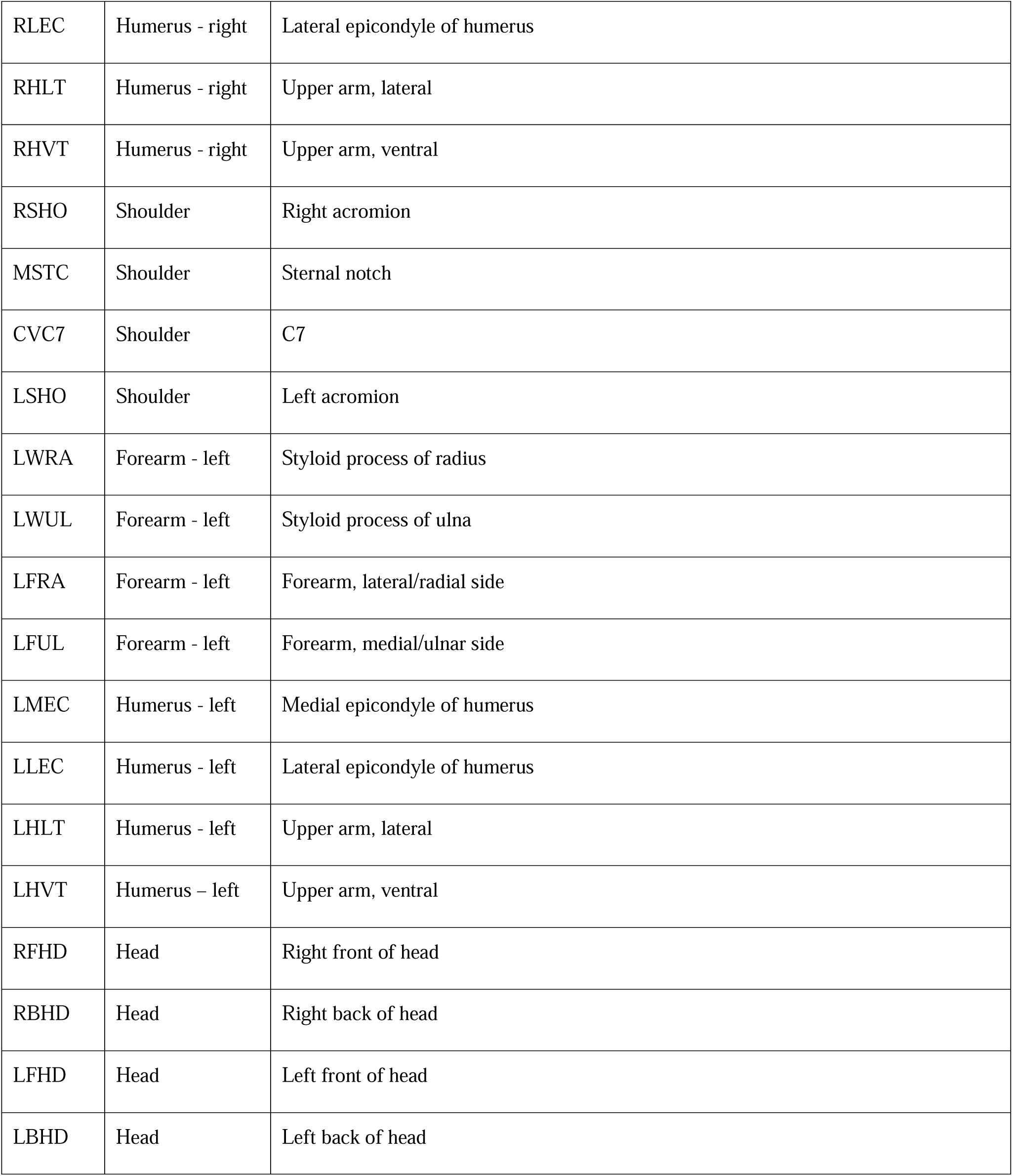
Locations of reflective markers.

**Supplementary Table 2.**
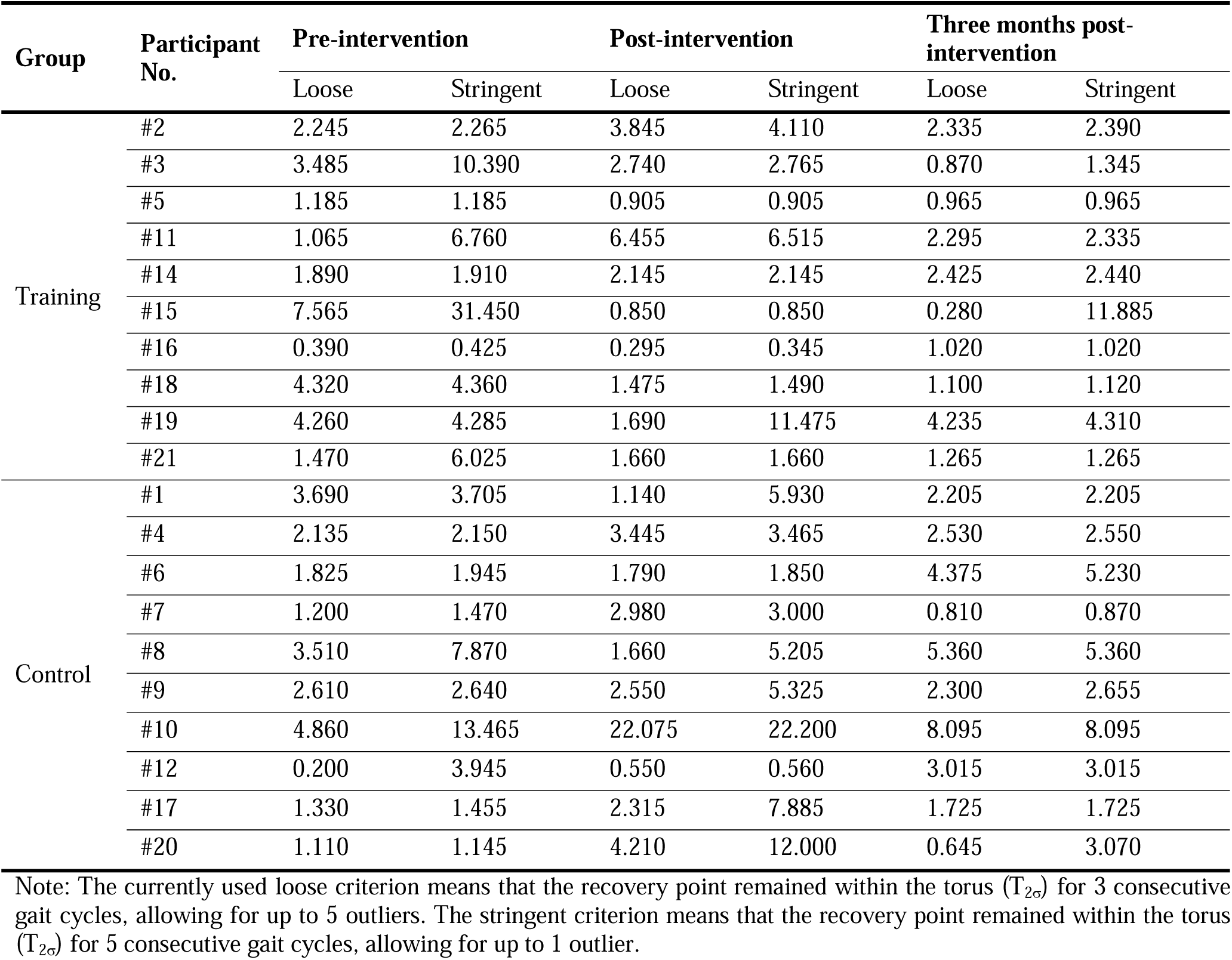
The recovery time values (s) based on two different criteria of identifying recovery points.

| Group | Participant No. | Pre-intervention |  | Post-intervention |  | Three months post-intervention |  |
| --- | --- | --- | --- | --- | --- | --- | --- |
|  |  | Loose | Stringent | Loose | Stringent | Loose | Stringent |
| Training | #2 | 2.245 | 2.265 | 3.845 | 4.110 | 2.335 | 2.390 |
|  | #3 | 3.485 | 10.390 | 2.740 | 2.765 | 0.870 | 1.345 |
|  | #5 | 1.185 | 1.185 | 0.905 | 0.905 | 0.965 | 0.965 |
|  | #11 | 1.065 | 6.760 | 6.455 | 6.515 | 2.295 | 2.335 |
|  | #14 | 1.890 | 1.910 | 2.145 | 2.145 | 2.425 | 2.440 |
|  | #15 | 7.565 | 31.450 | 0.850 | 0.850 | 0.280 | 11.885 |
|  | #16 | 0.390 | 0.425 | 0.295 | 0.345 | 1.020 | 1.020 |
|  | #18 | 4.320 | 4.360 | 1.475 | 1.490 | 1.100 | 1.120 |
|  | #19 | 4.260 | 4.285 | 1.690 | 11.475 | 4.235 | 4.310 |
|  | #21 | 1.470 | 6.025 | 1.660 | 1.660 | 1.265 | 1.265 |
| Control | #1 | 3.690 | 3.705 | 1.140 | 5.930 | 2.205 | 2.205 |
|  | #4 | 2.135 | 2.150 | 3.445 | 3.465 | 2.530 | 2.550 |
|  | #6 | 1.825 | 1.945 | 1.790 | 1.850 | 4.375 | 5.230 |
|  | #7 | 1.200 | 1.470 | 2.980 | 3.000 | 0.810 | 0.870 |
|  | #8 | 3.510 | 7.870 | 1.660 | 5.205 | 5.360 | 5.360 |
|  | #9 | 2.610 | 2.640 | 2.550 | 5.325 | 2.300 | 2.655 |
|  | #10 | 4.860 | 13.465 | 22.075 | 22.200 | 8.095 | 8.095 |
|  | #12 | 0.200 | 3.945 | 0.550 | 0.560 | 3.015 | 3.015 |
|  | #17 | 1.330 | 1.455 | 2.315 | 7.885 | 1.725 | 1.725 |
|  | #20 | 1.110 | 1.145 | 4.210 | 12.000 | 0.645 | 3.070 |

Note: The currently used loose criterion means that the recovery point remained within the torus ( $T_{2\sigma}$ ) for 3 consecutive gait cycles, allowing for up to 5 outliers. The stringent criterion means that the recovery point remained within the torus ( $T_{2\sigma}$ ) for 5 consecutive gait cycles, allowing for up to 1 outlier.

## References

1 Terroso, M., Rosa, N., Torres Marques, A. & Simoes, R. Physical consequences of falls in the elderly: a literature review from 1995 to 2010. Eur. Rev. Aging Phys. Act. 11, 51–59 (2014).

2 Mansfield, A., Wong, J. S., Bryce, J., Knorr, S. & Patterson, K. K. Does perturbation-based balance training prevent falls? Systematic review and meta-analysis of preliminary randomized controlled trials. Phys. Ther. 95, 700–709 (2015).

3 Salari, N., Darvishi, N., Ahmadipanah, M., Shohaimi, S. & Mohammadi, M. Global prevalence of falls in the older adults: a comprehensive systematic review and meta-analysis. J. Orthop. Surg. Res. 17, 334 (2022).

4 Niemann, S., Stürmer, Y., Meier, D. & Ellenberger, L. Status 2023: Statistics on non-occupational accidents and the level of safety in Switzerland. (2023).

5. World Health Organization. Falls, <https://www.who.int/news-room/fact-sheets/detail/falls#> (2021).

6 Duc, M. et al. Current practices of physiotherapists in Switzerland regarding fall risk-assessment for community-dwelling older adults: A national cross-sectional survey. F1000Research 11 (2022).

7 Sherrington, C. et al. Exercise for preventing falls in older people living in the community. Cochrane Database Syst. Rev. 1, Cd012424, doi:10.1002/14651858.CD012424.pub2 (2019).

8 Montero-Odasso, M. et al. World guidelines for falls prevention and management for older adults: a global initiative. Age and ageing 51, afac205 (2022).

9 Wang, Y. et al. Can a single session of treadmill-based slip training reduce daily life falls in community-dwelling older adults? A randomized controlled trial. Aging Clin. Exp. Res. 34, 1593–1602 (2022).

10 Luukinen, H. et al. Fracture risk associated with a fall according to type of fall among the elderly. Osteoporosis International 11, 631–634 (2000).

11 McCrum, C. et al. Perturbation-based balance training: Principles, mechanisms and implementation in clinical practice. Frontiers in sports and active living 4, 1015394 (2022).

12 Rogers, M. W. et al. Comparison of Lateral Perturbation-Induced Step Training and Hip Muscle Strengthening Exercise on Balance and Falls in Community-Dwelling Older Adults: A Blinded Randomized Controlled Trial. J. Gerontol. A Biol. Sci. Med. Sci. 76, e194–e202, doi:10.1093/gerona/glab017 (2021).

13 Nørgaard, J. E. et al. Effect of treadmill perturbation-based balance training on fall rates in community-dwelling older adults: a randomized clinical trial. JAMA network open 6, e238422–e238422 (2023).

14 Pai, Y.-C., Bhatt, T., Yang, F., Wang, E. & Kritchevsky, S. Perturbation training can reduce community-dwelling older adults’ annual fall risk: a randomized controlled trial. Journals of Gerontology Series A: Biomedical Sciences and Medical Sciences 69, 1586–1594 (2014).

15 Pai, Y.-C., Yang, F., Bhatt, T. & Wang, E. Learning from laboratory-induced falling: long-term motor retention among older adults. Age 36, 1367–1376 (2014).

16 Wang, Y. et al. Can treadmill-slip perturbation training reduce immediate risk of over-ground-slip induced fall among community-dwelling older adults? J. Biomech. 84, 58–66 (2019).

17 Wang, Y., Wang, S., Lee, A., Pai, Y.-C. & Bhatt, T. Treadmill-gait slip training in community-dwelling older adults: mechanisms of immediate adaptation for a progressive ascending-mixed-intensity protocol. Exp. Brain Res. 237, 2305–2317 (2019).

18 Lee, A., Bhatt, T., Liu, X., Wang, Y. & Pai, Y.-C. Can higher training practice dosage with treadmill slip-perturbation necessarily reduce risk of falls following overground slip? Gait Posture 61, 387–392 (2018).

19 Liu, X. et al. The retention of fall-resisting behavior derived from treadmill slip-perturbation training in community-dwelling older adults. Geroscience 43, 913–926 (2021).

20 Wang, Y., Wang, S., Bolton, R., Kaur, T. & Bhatt, T. Effects of task-specific obstacle-induced trip-perturbation training: proactive and reactive adaptation to reduce fall-risk in community-dwelling older adults. Aging Clin. Exp. Res. 32, 893–905 (2020).

21 Bhatt, T., Wang, Y., Wang, S. & Kannan, L. Perturbation training for fall-risk reduction in healthy older adults: Interference and generalization to opposing novel perturbations post intervention. Frontiers in sports and active living 3, 697169 (2021).

22 Song, P. Y., Sturnieks, D. L., Davis, M. K., Lord, S. R. & Okubo, Y. Perturbation-based balance training using repeated trips on a walkway vs. belt accelerations on a treadmill: a cross-over randomised controlled trial in community-dwelling older adults. Frontiers in sports and active living 3, 702320 (2021).

23 Watson, F. et al. Use of the margin of stability to quantify stability in pathologic gait–a qualitative systematic review. BMC musculoskeletal disorders 22, 1–29 (2021).

24 Karamanidis, K., Epro, G., McCrum, C. & König, M. Improving trip-and slip-resisting skills in older people: perturbation dose matters. Exercise and sport sciences reviews 48, 40–47 (2020).

25 McCrum, C., Karamanidis, K., Willems, P., Zijlstra, W. & Meijer, K. Retention, savings and interlimb transfer of reactive gait adaptations in humans following unexpected perturbations. Communications biology 1, 230 (2018).

26 Debelle, H., Harkness-Armstrong, C., Hadwin, K., Maganaris, C. N. & O’Brien, T. D. Recovery from a forward falling slip: measurement of dynamic stability and strength requirements using a split-belt instrumented treadmill. Frontiers in sports and active living 2, 82 (2020).

27 Stergiou, N. Nonlinear analysis for human movement variability. (CRC press, 2018).

28 Ravi, D. K. et al. Rhythmic auditory stimuli modulate movement recovery in response to perturbation during locomotion. J. Exp. Biol. 224, jeb237073 (2021).

29 Ravi, D. K., Heimhofer, C. C., Taylor, W. R. & Singh, N. B. Adapting footfall rhythmicity to auditory perturbations affects resilience of locomotor behavior: A proof-of-concept study. Front. Neurosci. 15, 678965 (2021).

30 O’Connor, C. M., Thorpe, S. K., O’Malley, M. J. & Vaughan, C. L. Automatic detection of gait events using kinematic data. Gait Posture 25, 469–474 (2007).

31 Yang, F. & Pai, Y.-C. Can sacral marker approximate center of mass during gait and slip-fall recovery among community-dwelling older adults? J. Biomech. 47, 3807–3812 (2014).

32 Hof, A. L., Gazendam, M. & Sinke, W. The condition for dynamic stability. J. Biomech. 38, 1–8 (2005).

33 Curtze, C., Buurke, T. J. & McCrum, C. Notes on the margin of stability. J. Biomech. 166, 112045 (2024).

34 McCrum, C. et al. Deficient recovery response and adaptive feedback potential in dynamic gait stability in unilateral peripheral vestibular disorder patients. Physiological reports 2, e12222 (2014).

35 Wurdeman, S. R. in Nonlinear analysis for human movement variability 55–82 (CRC Press, 2018).

36 Podsiadlo, D. & Richardson, S. The timed “Up & Go”: a test of basic functional mobility for frail elderly persons. J. Am. Geriatr. Soc. 39, 142–148 (1991).

37 Barker, K. L. et al. Physiotherapy Rehabilitation for Osteoporotic Vertebral Fracture (PROVE): study protocol for a randomised controlled trial. Trials 15, 1–11 (2014).

38 Valkovič, P., Brožová, H., Bötzel, K., Růžička, E. & Benetin, J. Push and release test predicts better Parkinson fallers and nonfallers than the pull test: comparison in OFF and ON medication states. Movement disorders: official journal of the Movement Disorder Society 23, 1453–1457 (2008).

39 Cohen, J. Statistical power analysis for the behavioral sciences. (routledge, 2013).

40 Gerards, M. H. et al. Adaptability to balance perturbations during walking as a potential marker of falls history in older adults. Frontiers in sports and active living 3, 132 (2021).

41 Zhu, R. T.-L. et al. How Does Lower Limb Respond to Unexpected Balance Perturbations? New Insights from Synchronized Human Kinetics, Kinematics, Muscle Electromyography (EMG) and Mechanomyography (MMG) Data. Biosensors 12, 430 (2022).

42 Tong, C. Y., et al. Muscular and Kinematic Responses to Unexpected Translational Balance Perturbation: A Pilot Study in Healthy Young Adults. Bioengineering 10 (2023).

43 Kasahara, S. & Saito, H. Mechanisms of postural control in older adults based on surface electromyography data. Human Movement Science 78, 102803, 10.1016/j.humov.2021.102803 (2021).

44 Zhu, R. T.-L. et al. Older Fallers and Non-fallers’ Neuromuscular and Kinematic Alterations in Reactive Balance Control: Indicators of Balance Decline or Compensation? (2024).

45 Brüll, L., Hezel, N., Arampatzis, A. & Schwenk, M. Comparing the effects of two perturbation-based balance training paradigms in fall-prone older adults: a randomized controlled trial. Gerontology 69, 910–922 (2023).

46 Mansfield, A., Peters, A. L., Liu, B. A. & Maki, B. E. Effect of a perturbation-based balance training program on compensatory stepping and grasping reactions in older adults: a randomized controlled trial. Phys. Ther. 90, 476–491 (2010).

47 Lurie, J. D. et al. Surface perturbation training to prevent falls in older adults: a highly pragmatic, randomized controlled trial. Phys. Ther. 100, 1153–1162 (2020).

48 Aviles, J. et al. Comparison of treadmill trip-like training versus Tai Chi to improve reactive balance among independent older adult residents of senior housing: a pilot controlled trial. The Journals of Gerontology: Series A 74, 1497–1503 (2019).

49 Allin, L. J. et al. Perturbation-based balance training targeting both slip-and trip-induced falls among older adults: a randomized controlled trial. BMC Geriatr. 20, 1–13 (2020).

50 Niewolak, K. et al. Assessment of postural control and gait in late post-stroke patients subjected to treadmill training with controlled body balance perturbations: pilot study. *Issues of Rehabilitation, Orthopaedics*, Neurophysiology and Sport Promotion-IRONS 39 (2022).

51. Bhaskar, B., Jimshad, T. & Solomen, S. Efficacy of perturbation training in improving balance and function in the management of knee osteoarthritis. (2019).

52. Todd, C. & Skelton, D. What are the main risk factors for falls amongst older people and what are the most effective interventions to prevent these falls?, (World Health Organization. Regional Office for Europe, 2004).

